# Multi-stage models for the failure of complex systems, cascading disasters, and the onset of disease

**DOI:** 10.1101/476242

**Authors:** Anthony J. Webster

## Abstract

Complex systems can fail through different routes, often progressing through a series of (rate-limiting) steps and modified by environmental exposures. The onset of disease, cancer in particular, is no different. Multi-stage models provide a simple but very general mathematical framework for studying the failure of complex systems, or equivalently, the onset of disease. They include the Armitage-Doll multi-stage cancer model as a particular case, and have potential to provide new insights into how failures and disease, arise and progress. A method described by E.T. Jaynes is developed to provide an analytical solution for a large class of these models, and highlights connections between the convolution of Laplace transforms, sums of random variables, and Schwinger/Feynman parameterisations. Examples include: exact solutions to the Armitage-Doll model, the sum of Gamma-distributed variables with integer-valued shape parameters, a clonal-growth cancer model, and a model for cascading disasters. Applications and limitations of the approach are discussed in the context of recent cancer research. The model is sufficiently general to be used in many contexts, such as engineering, project management, disease progression, and disaster risk for example, allowing the estimation of failure rates in complex systems and projects. The intended result is a mathematical toolkit for applying multi-stage models to the study of failure rates in complex systems and to the onset of disease, cancer in particular.

## 1 Introduction

Complex systems such as a car can fail through many different routes, often requiring a sequence or combination of events for a component to fail. The same can be true for human disease, cancer in particular [1, 2, 3]. For example, cancer can arise through a sequence of steps such as genetic mutations, each of which must occur prior to cancer [4, 5, 6, 7, 8]. The considerable genetic variation between otherwise similar cancers [9, 10], suggests that similar cancers might arise through a variety of different paths.

Multi-stage models describe how systems can fail through one or more possible routes. They are sometimes described as “multi-step” or “multi-hit” models [11, 12], because each route typically requires failure of one or more sequential or non-sequential steps. Here we show that the model is easy to conceptualise and derive, and that many specific examples have analytical solutions or approximations, making it ideally suited to the construction of biologically- or physically-motivated models for the incidence of events such as diseases, disasters, or mechanical failures. A method described by E.T. Jaynes [13] generalises to give an exact analytical formula for the sums of random variables needed to evaluate the sequential model. This is evaluated for specific cases. Moolgavkar’s exact solution [14] to the Armitage-Doll multistage cancer model is one example that is derived surprisingly easily, and is easily modified. The approach described here can incorporate simple models for a clonal expansion prior to cancer detection [5, 6, 7], but as discussed in Sections 8 and 9, it may not be able to describe evolutionary competition or cancer-evolution in a changing micro-environment without additional modification. More generally, it is hoped that the mathematical framework can be used in a broad range of applications, including the modelling of other diseases [15, 16, 17, 18]. One example we briefly describe in Section 8 is modelling of “cascading disasters” [19], where each disaster can substantially modify the risk of subsequent (possibly different) disasters.

Conventional notation is used [20], with: probability densities *f*(*t*), cumulative probability distributions 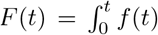, a survival function *S*(*t*) = 1 − *F*(*t*), hazard function *h*(*t*) = *f*(*t*)/*S*(*t*), and cumulative hazard function 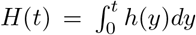. Noting that *f*(*t*) = −*dS*/*dt*, it is easily seen that 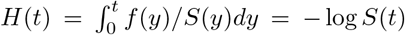, *h*(*t*) = −*d*log *S*(*t*)/*dt*, and 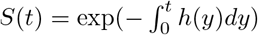.

## 2 Failure by multiple possible routes

Imagine that we can enumerate all possible routes 1 to *n* by which a failure can occur (Fig 1). The probability of surviving the *i*th of these routes after time *t* is *S_i_*(*t*), and consequently the probability of surviving all of these possible routes to failure *S*(*t*) is,

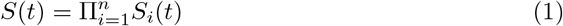

or in terms of cumulative hazard functions with *S_i_*(*t*) = *e*^−*H_i_*(*t*)^,

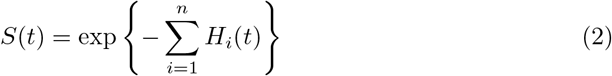

**Figure 1:**
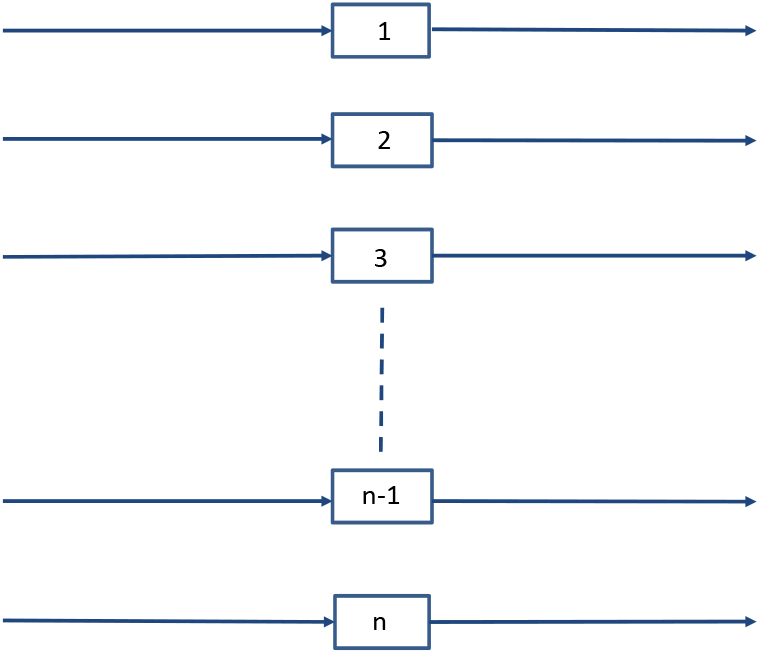
In a complex system, failure can occur through many different routes (Eq. 1).

The system’s hazard rate for failure by any of the routes is,

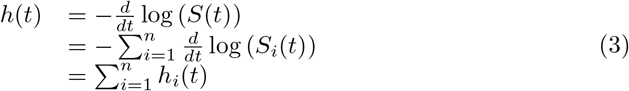

and 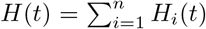. In words, if failure can occur by any of *n* possible routes, the overall hazard of failure equals the sum of the hazard of failure by all the individual routes.

A few notes on Eq. 2 and its application to cancer modelling. Firstly, if the *s*th route to failure is much more likely than the others, with *H_s_* ≫ *H_j_* for *s* ≠ *j*, then *S*(*t*) = exp {−*H_s_*(*t*) + (1 + *O* (Σ_*i*≠*s*_ *H_i_*/*H_s_*))} ≃ exp {−*H_s_*(*t*)}, which could represent the most likely sequence of mutations in a cancer model for example. Due to different manufacturing processes, genetic backgrounds, chance processes or exposures (e.g. prior to adulthood), this most probable route to failure could differ between individuals. Secondly, the stem cell cancer model assumes that cancer can occur through any of *n_s_* equivalent stem cells in a tissue, for which Eq. 2 is modified to, 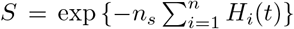. So a greater number of stem cells is expected to increase cancer risk, as is observed [21, 22]. Thirdly, most cancers are sufficiently rare that *S* ~ 1. As a consequence, many cancer models (implicity or explicitly) assume 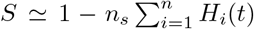 and 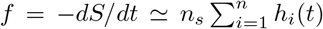, a limit emphasised in the Appendix of Moolgavkar [14].

## 3 Failure requiring *m* independent events

Often failure by a particular path will require more than one failure to occur independently. Consider firstly when there are *m_i_* steps to failure, and the order of failure is unimportant (Fig 2). The probability of surviving failure by the *i*th route, *S_i_*(*t*) is,

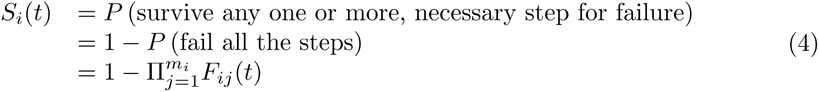

where *F_ij_*(*t*) is the cumulative probability distribution for failure of the *j*th step on the *i*th route within time *t*. Writing *S_ij_*(*t*) = 1 − *F_ij_*(*t*), this can alternately be written as,

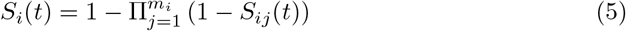

**Figure 2:**
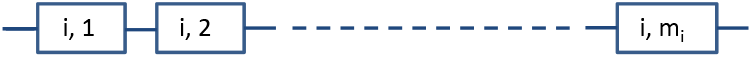
Failure by the ith path at time t requires m_i_ independent failures to occur in any order, with the last failure at time t (Eq. 5).

## 4 Relation to recent multi-stage cancer models

It may be helpful to explain how Eqs. 1 and 4 are used in recently described multi-stage cancer models [23, 24, 25]. If we take a rate of mutations *μ_j_* per cell division for each of the rate-limiting mutational steps 1 to *j*, and *d_i_* divisions of cell *i*, then the probability of a stem cell surviving without the jth rate limiting mutation is *S_ij_* = (1 − *μ_j_*)^*d_i_*^. Similarly, the probability of a given stem cell having mutation *j* is *F_ij_* = 1 − (1 − *μ_j_*)^*d_i_*^. This is the solution of Zhang et al. [24] to the recursive formula of Wu et al. [23] (see Appendix of Zhang et al. [24] for details). Using Eq. 4, the survival of the *i*th stem cell is described by,

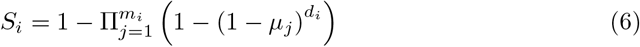

Now assuming all *n* stem cells are equivalent and have equal rates *μ_i_* = *μ_j_* for all *i, j*, and consider only one path to cancer with *m* mutational steps, then,

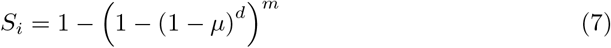

and,

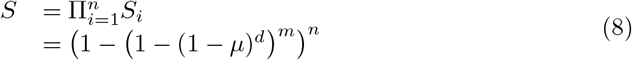

The probability of cancer within *m* divisions, often referred to as “theoretical lifetime intrinsic cancer risk”, is,

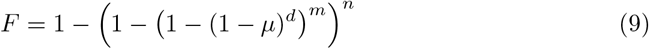

This is the equation derived by Calabrese and Shibata [25], and that Zhang found as the solution to the model of Wu et al [24, 23].

Therefore, in addition to the models of Wu and Calabrese being equivalent cancer models needing m mutational steps, the models also assume that the order of the steps is not important. This differs from the original Armitage-Doll model that considered a sequential set of rate-limiting steps, and was exactly solved by Moolgavkar [14]. Eqs. 8 and 9 are equivalent to assuming: (i) equivalent stem cells, (ii) a single path to cancer, (iii) equivalent divisions per stem cell, and, (iv) equivalent mutation rates for all steps.

Despite the differences in modelling assumptions for Eq. 9 and the Armitage-Doll model, their predictions can be quantitatively similar. To see this, use the Armitage-Doll approximation of *μd* ≪ 1, to expand,

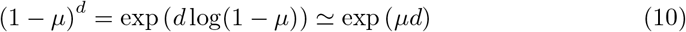

If cell divisions are approximately uniform in time, then we can replace *μd* with *μt*, with *μ* now a rate per unit time. Then expanding exp(−*μt*) ≃ 1 − *μt*, gives,

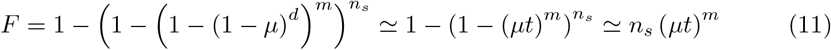

The incidence rate *h* = *f*/*S* is then *h* ≃ *n_s_μ^m^t*^*m*−1^, the same as the original (approximate) Armitage-Doll solution [2]. This approximate solution is expected to become inaccurate at sufficiently long times.

An equivalent expression to Eq. 8 was known to Armitage, Doll, and Pike since at least 1965 [26], as was its limiting behaviour for large *n*. The authors [26] emphasised that many different forms for the *F_i_*(*t_i_*) could produce approximately the same observed *F*(*t*), especially for large *n*, with the behaviour of *F*(*t*) being dominated by the small *t* behaviour of *F_i_*(*t*). As a result, for sufficiently small times power-law behaviour for *F*(*t*) is likely, and if longer times were observable then an extreme value distribution would be expected [26, 27, 4]. However the power-law approximation can fail for important cases with extra rate-limiting steps such as a clonal expansion [5, 6, 7]. It seems likely that a model that includes clonal expansion and cancer detection is needed for cancer modelling, but the power law approximation could be used for all but the penultimate step, for example. A general methodology that includes this approach is described next, and examples are given in the subsequent section 6. The results and examples of sections 5 and 6 are intended to have a broad range of applications.

## 5 Failure requiring *m* sequential steps

Some failures require a *sequence* of independent events to occur, each following the one before (Fig 3). A well-known example is the Armitage-Doll multistage cancer model, that requires a sequence of *m* mutations (failures), that each occur with a different constant rate. The probability density for failure time is the pdf for a sum of the *m* independent times *t_j_* to failure at each step in the sequence, each of which may have a different probability density function *f_j_*(*t_j_*). A general method for evaluating the probability density is outlined below, adapting a method described by Jaynes [13] (page 569).

**Figure 3:**
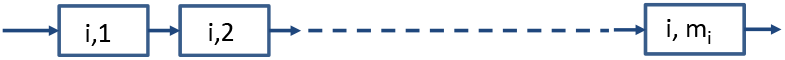
Failure by the ith path at time t requires an ordered sequence of failures, with the last failure at time t (Eqs. 16 and 18).

Take *T_i_* ~ *f_i_*(*t_i_*) as random variables. Then use marginalisation to write 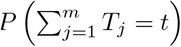 in terms of 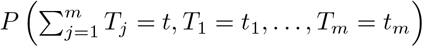, where (*A, B, C*) is read as “*A* and *B* and *C*”, and expand using the product rule *P*(*A, B*) = *P*(*A*|*B*)*P*(*B*),

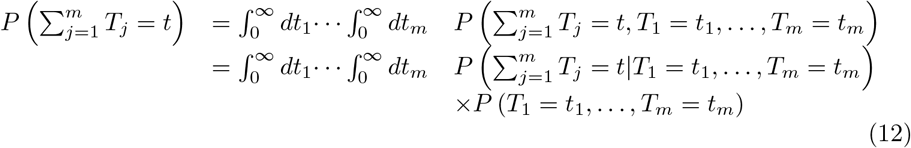

Noting that 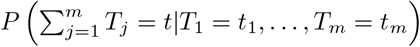 is zero for 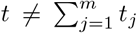 and 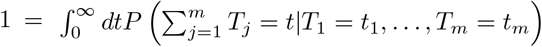, indicates that it is identical to a Dirac delta function 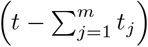. For independent events 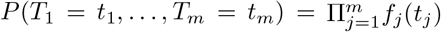 where *f_j_*(*t_j_*) ≡ *P_j_*(*T_j_* = *t_j_*). Writing 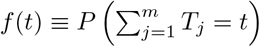, then gives,

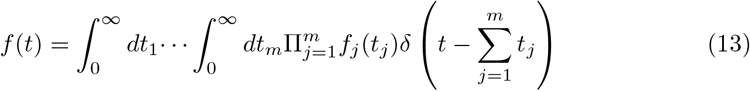

To evaluate the integrals, take the Laplace transform with respect to *t*, to give,

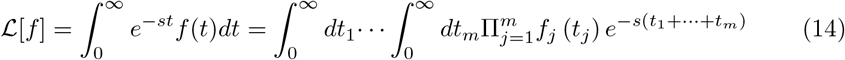

This factorises as,

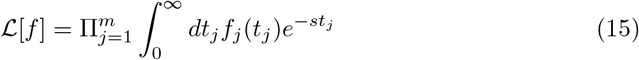

Giving a general analytical solution as,

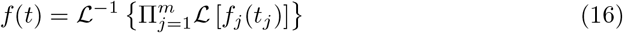

where 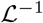 is the inverse Laplace transform, and 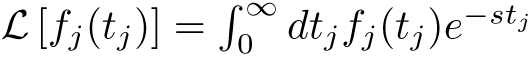 with the same variable s for each value of *j*. Eq. 15 is similar to the relationship between moment generating functions 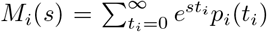 of discrete probability distributions *p_i_*(*t_i_*), and the moment generating function *M*(*s*) for 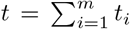, that has,

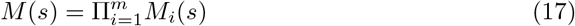

whose derivation is analogous to Eq. 17 but with integrals replaced by sums. The survival and hazard functions for *f*(*t*) can be obtained from Eq. 16 in the usual way. For example,

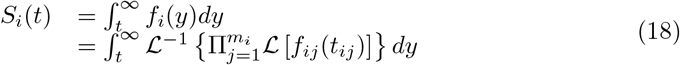

that can be used in combination with Eq. 1. A number of valuable results are easy to evaluate using Eq. 16, as is illustrated in the next section.

A useful related result is,

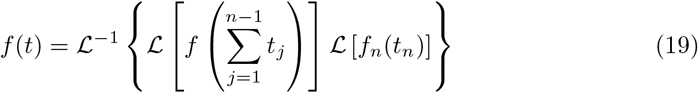

that can be inferred from Eq. 16 with *m* = 2,

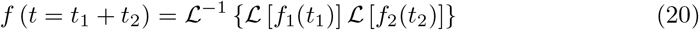

by replacing *f*_1_(*t*_1_) with 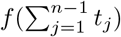 and *f*_2_(*t*_2_) with *f_n_*(*t_n_*). Eq. 20 can be solved using the convolution theorem for Laplace transforms, that gives,

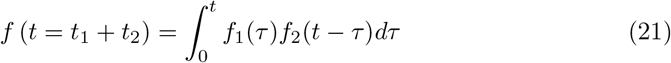

which is sometimes easier to evaluate than two Laplace transforms and their inverse. In general, solutions can be presented in terms of multiple convolutions if it is preferable to do so. Eqs. 19 and 21 are particularly useful for combining a known solution for the sum of (*n* − 1) samples such as for cancer initiation, with a differently distributed *n*th sample, such as the waiting time to detect a growing cancer. A final remark applies to the sum of random variables whose domain extends from −∞ to ∞, as opposed to the range 0 to ∞ considered so far. In that case an analogous calculation using a Fourier transform with respect to *t* in Eq. 13 leads to analogous results in terms of Fourier transforms, with 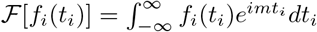 in place of Laplace transforms, resulting in,

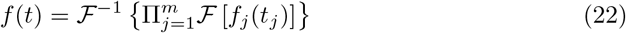

Eq. 22 is mentioned for completeness, but is not used here.

A general solution to Eq. 16 can be given in terms of definite integrals, with,

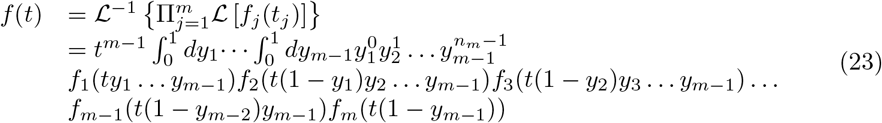

This can sometimes be easier to evaluate or approximate than Eq. 16. A derivation is given in the Supporting Information (S1 Appendix). Eq. 23 allows a generalised Schwinger/Feynman parameterisation [28] to be derived. Writing 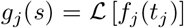 and taking the Laplace transform of both sides of Eq. 23, gives,

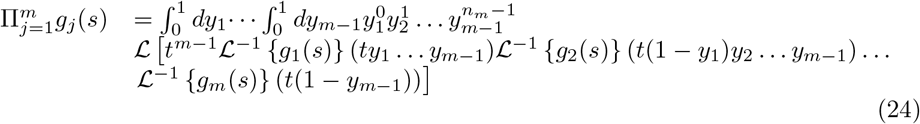

which includes some well known Schwinger/Feynman parameterisations as special cases. This is discussed further in the Supporting Information (S1 Appendix).

## 6 Modelling sequential events - examples

In the following examples we consider the time 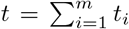 for a sequence of events, with possibly different distributions *f_i_*(*t_i_*) for the time between the (*i* − 1)th and *i*th event. Some of the results are well-known but not usually presented this way, others are new or poorly known. We will use the Laplace transforms (and their inverses), of,

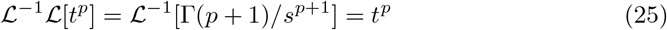

and

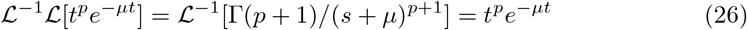

### Sums of gamma distributed samples (equal rates)

Using Eq. 16, the sum of *m* gamma distributed variables with equal rate parameters *μ*, and 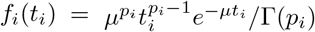, are distributed as,

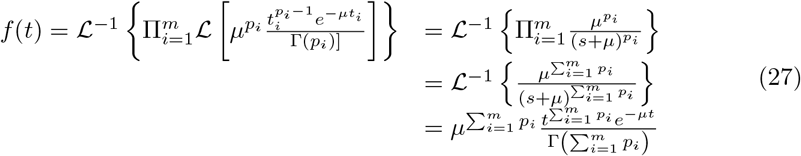

For a sum of m exponentially distributed variables with {*p_i_* = 1}, this simplifies to *f*(*t*) = *μ^m^t*^*m*−1^*e*^−*μt*^/Γ(*m*), a Gamma distribution.

### Power law approximations

For many situations such as most diseases, you are unlikely to get any particular disease during your lifetime. In those cases the probability of survival over a lifetime is close to 1, and the probability density function *f_i_* = *h_i_*/*S_i_*, can be approximated by *f_i_* ≃ *h_i_*, that in turn can often be approximated by a power of time with 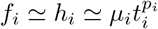. Then we have,

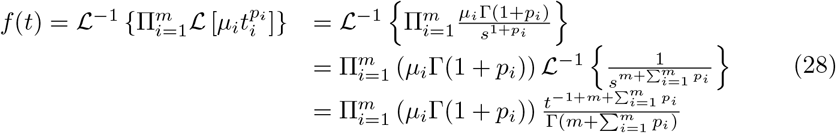

### The Armitage-Doll model

A well known example of this approximation Eq. 28, is (implicitly) in the original approximate solution to the Armitage-Doll multi-stage cancer model. Taking a constant hazard at each step, and approximating *f_i_* ≃ *h_i_* = *μ_i_*, then Eq. 28 gives,

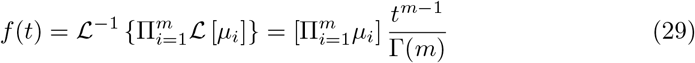

as was used in the original Armitage-Doll paper. Note that an equivalent time-dependence can be produced by a different combination of hazard functions with 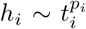 and 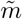 steps, provided 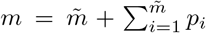. For example, if *m* = 6, there could be 3 steps with *p* = 1, or 2 steps with *p* = 2, or 3 steps with *p*_1_ = 0, *p*_2_ = 1, and *p*_3_ = 2, or some more complex combination. If the full pdfs are modelled at each step as opposed to their polynomial approximation, then this flexibility is reduced, as is the case for Moolgavkar’s exact solution to the Armitage-Doll model that is described next.

### Moolgavkar’s exact solution to the Armitage-Doll model

Moolgavkar’s exact solution to the Armitage-Doll model is the solution of,

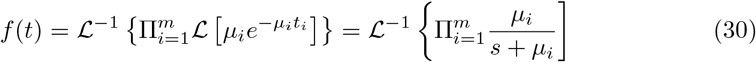

For example, if *n* = 3 then,

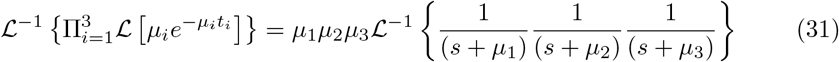

Using partial fractions, we can write,

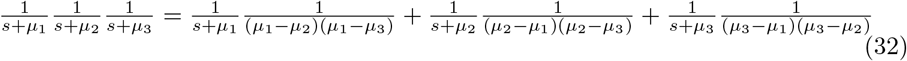

Allowing the inverse Laplace transforms to be easily evaluated, giving,

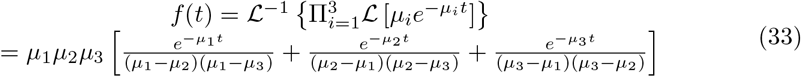

Note that the result is independent of the order of sequential events, but unlike the approximate solution to the Armitage Doll model [2], the exact solution allows less variability in the underlying models that can produce it. Also note that the leading order terms of an expansion in t cancel exactly, to give identical leading-order behaviour as for a power-law approximation (with *p* = 0)

A general solution can be formed using a Schwinger/Feynman parameterisation [28] of,

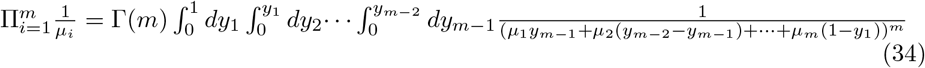

Replacing *μ_i_* with *s* + *μ_i_* in Eq. 34, then we can write Eq. 30 as,

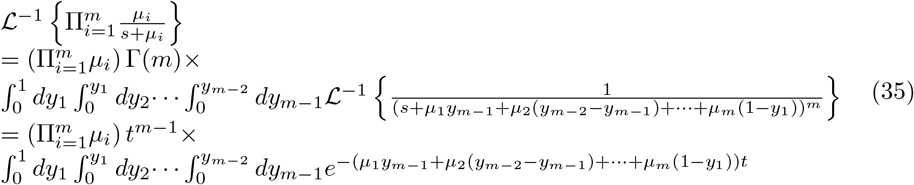

(which is simpler, but equivalent in effect, to repeatedly using the convolution formula). Completing the integrals will generate Moolgavkar’s solution for a given value of *m*. For example, taking *m* = 3 and integrating once gives,

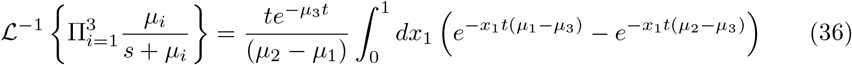

Integrating a second time, and simplifying, gives Eq. 33. The relationships between Schwinger/Feynman parameterisations, Laplace transforms, and the convolution theorem are discussed further in the Supplementary Information (S1 Appendix).

Moolgavkar [14] used induction to provide an explicit formula for *f*(*t*), with,

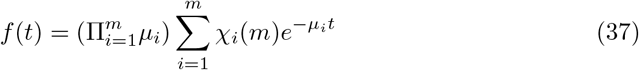

where,

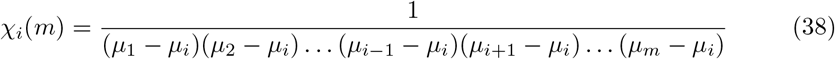

For small times the terms in a Taylor expansion of Eq. 37 cancel exactly, so that 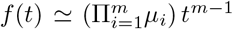, as expected. This feature could be useful for approximating a normalised function when the early-time behaviour approximates an integer power of time. Further uses of Moolgavkar’s solution are discussed next.

### Sums of gamma distributed samples (with different rates)

A useful mathematical result can be found by combining the Laplace transform of Moolgavkar’s solution Eq. 37 for 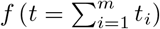 with Eq. 30, to give an explicit formula for a partial fraction decomposition of the product 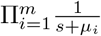, as,

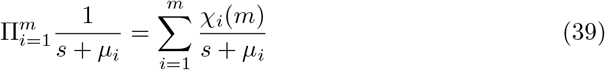

This can be useful in various contexts. For example, consider *m* Gamma distributions 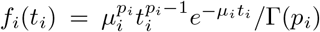 with different integer-valued shape parameters *p_i_*, and 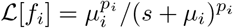. Eq. 16 gives 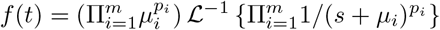, so firstly use the integer-valued property of {*p_i_*} to write,

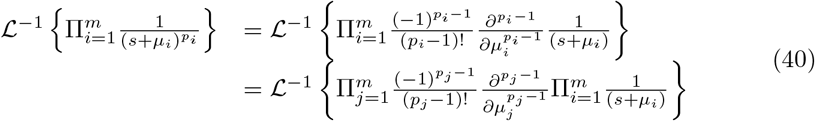

where the product of differential operators can be taken outside the product of Laplace transforms because *∂*/*∂μ_i_*(1/(*s* + *μ_j_*)) is zero for *i* ≠ *j*. Using Eq. 39 we can replace the product of Laplace transforms with a sum, giving,

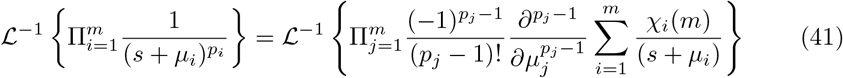

The Laplace transform has now been simplified to a sum of terms in 1/(*s* + *μ_i_*), whose inverse Laplace transforms are easy to evaluate. Taking the inverse Laplace transform 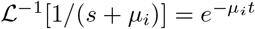, and including the product 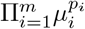, gives,

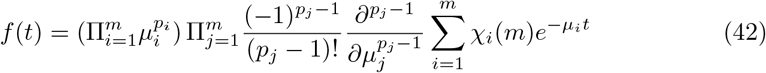

as a general solution for sums of Gamma distributed samples with integer-valued shape parameters *p_i_* (and arbitrary rate parameters *μ_i_*). Eq. 42 is most easily evaluated with a symbolic algebra package.

If *p_i_* = *p* are equal, then Eq. 42 may be simplified further by writing,

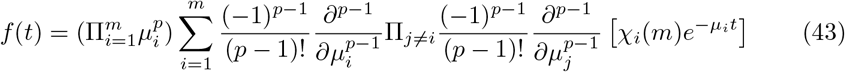

and noting that,

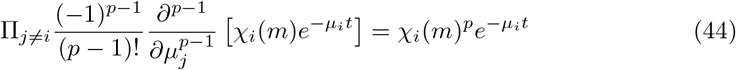

because for *j* = *i* there is exactly one factor 1/(*μ_j_* − *μ_i_*) in *χ_i_*(*m*). This leaves,

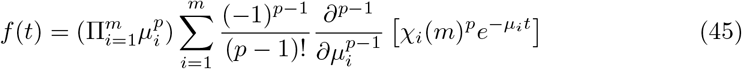

for sums of Gamma distributed samples with the same integer-valued shape parameter *p* (and arbitrary rate parameters *μ_i_*).

For example, if *p* = 1 then Eq. 45 becomes Moolgavkar’s Eq. 37. Alternatively, if e.g. *p* = 2, then we have,

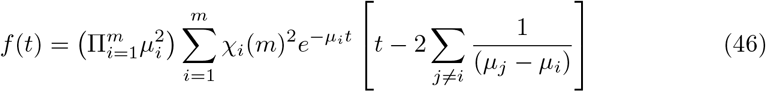

for the sum of Gamma distributions with shape parameters *p* = 2 and arbitrary rate parameters, and *χ_i_*(*m*) as defined in Eq. 38. If we also let e.g. *m* = 2, *μ*_2_ = *μ*_1_ + *ϵ*, and *ϵ* → 0, then Eq. 46 tends to 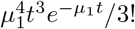, for the sum of two Gamma distributed variables with rate *μ*_1_ and *p* = 2, in agreement with Eq. 27.

### Sums of samples with different distributions

An advantage of the method described above, is that it is often easy to calculate pdfs for sums of differently distributed samples. For the first example, consider two samples from the same (or very similar) exponential distribution, and a third from a different exponential distribution. The result can be obtained by writing *μ*_3_ = *μ*_2_ + *ϵ* in Eq. 33, and letting *ϵ* → 0. Considering the terms involving exponents of *μ*_2_ and *μ*_3_,

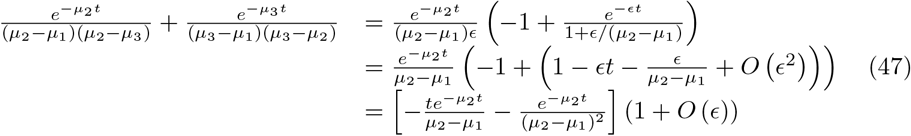

Using Eq. 33 and letting *ϵ* → 0, gives,

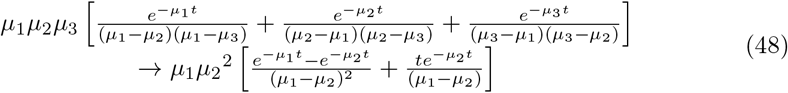

for the sum of three exponentially distributed variables, when exactly two have the same rate. Taking *μ*_2_ = *μ*_1_ + *ϵ* and letting *ϵ* → 0 in Eq. 48, gives a Gamma distribution 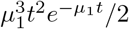, as it should for the sum of three exponentially distributed variables with equal rates (see Eq. 27 with {*p_i_* = 0}). More generally, it can be seen that a sum of exponentially distributed samples with different rates, smoothly approximate a gamma distribution as the rates become increasingly similar, as expected from Eq. 27.

### Failure involving a combination of sequential and non-sequential steps

If a path to failure involves a combination of sequential and non-sequential steps, then the necessary set of sequential steps can be considered as one of the non-sequential steps, with overall survival given by Eq. 1 and the survival for any sequential set of steps calculated from Eq. 18 (Fig 4).

**Figure 4:**
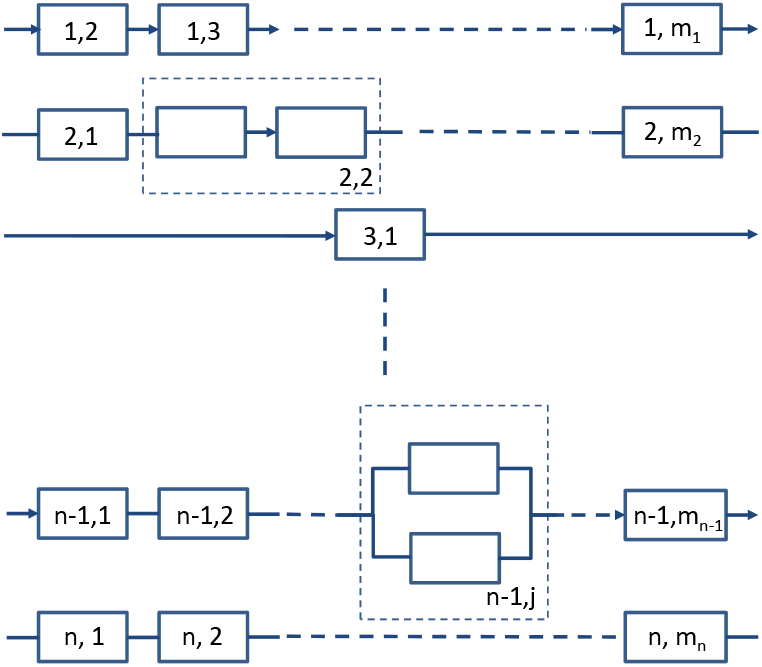
Overall failure risk can be modelled as sequential steps (e.g. (1,1) to (1,m_1_) using Eq. 5), and non-sequential steps (e.g. (n,1) to (n,m_n_) using Eq. 16), that may be dependent on each other (e.g. Eq. 55). For the purposes of modelling, a sequence of dependent or multiple routes can be regarded as a single step (e.g. (2,2) or (n − 1,j)).

## 7 Clonal-expansion cancer models

Clonal expansion is thought to be an essential element of cancer progression [29], and can modify the timing of cancer onset and detection [5, 6, 7, 32, 31, 30]. The growing number of cells at risk increases the probability of the next step in a sequence of mutations occurring, and if already cancerous, then it increases the likelihood of detection.

Some cancer models have a clonal expansion of cells as a rate-limiting step [5, 6, 7]. For example, Michor et al. [6] modelled clonal expansion of myeloid leukemia as logistic growth, with the likelihood of cancer detection (the hazard function), being proportional to the number of cancer cells. This gives a survival function for cancer detection of,

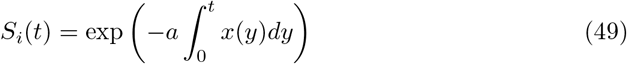

where,

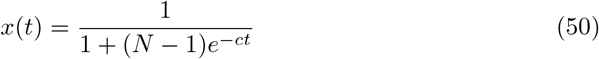

*a, c*, are rate constants, and *N* is the total number of cells prior to cancer initiation. Noting that 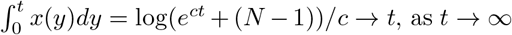, as *t* → ∞ and *x*(*t*) ∞ 1, then the tail of the survival curve falls exponentially towards zero with time.

Alternatively, we might expect the likelihood of cancer being diagnosed to continue to increase with time since the cancer is initiated. For example, a hazard function that is linear in time would give a Weibull distribution with *S*(*t*) = *e*^−*at*^2^^. It is unlikely that either this or the logistic model would be an equally good description for the detection of all cancers, although they may both be an improvement on a model without either. Both models have a single peak, but differ in the tail of their distribution, that falls as ~ *e*^−*act*^ for the logistic model and ~ *e*^−*at*^2^^ for the Weibull model. Qualitatively, we might expect a delay between cancer initiation and the possibility of diagnosis, and diagnosis to occur almost inevitably within a reasonable time-period. Therefore a Weibull or Gamma distributed time to diagnosis may be reasonable for many cancers, with the shorter tail of the Weibull distribution making it more suitable approximation for cancers whose diagnosis is almost inevitable. (The possibility of misdiagnosis or death by another cause is not considered here.)

For example, noting that Moolgavkar’s solution is a linear combination of exponential distributions, to combine it with a Weibull distribution for cancer detection 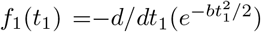, we can consider a single exponential term at a time. Taking *f*_2_(*t*_2_) = *ae*^−*at*^2^^, and using the convolution formula Eq. 21, we get,

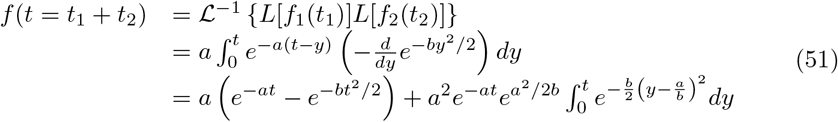

where we integrated by parts to get the last line. This may be written as,

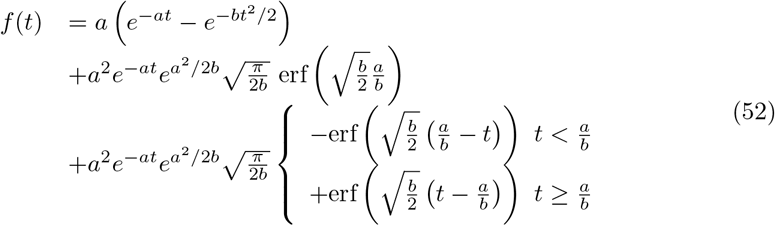

with 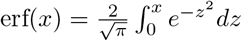. Similarly for a Gamma distribution with *f*_1_ = *b^p^t*^*p*−1^*e*^−*bt*^/Γ(*p*) and an exponential, *f_2_*(*t*_2_) = *ae*^−*at*^, then assuming *b* > *a*,

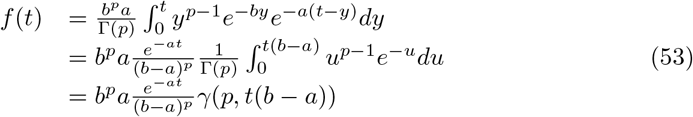

where *γ*(*p, t*(*b* − *a*)) is the normalised lower incomplete Gamma function, which is available in most computational mathematics and statistics packages. If *a* > *b* then f1 and f2 must be exchanged and the result is most easily evaluated numerically.

## 8 Cascading failures with dependent sequences of events

Now consider non-independent failures, where the failure of A changes the probability of a failure in B or C. In general, if the paths to failure are not independent of each other then the situation cannot be described by Eq. 1. Benjamin Cairns suggested exploring the following example - if step 1 of A prevents step 1 of B and vice-versa, then only one path can be followed. If the first step occurs at time *t*_1_, the pdf for failure at time *t* is: *f*(*t*) = *S_A_*(*t*_1_)*f_B_*(*t*) + *S_B_*(*t*_1_)*f_A_*(*t*), where *f_A_*(*t*) and *f_B_*(*t*) are the pdfs for path A and B if they were independent. This differs from Eq. 1 that has, *f*(*t*) = −*dS*/*dt* = *S_A_*(*t*)*f_B_*(*t*) + *S_B_*(*t*)*f_A_*(*t*), that is independent of *t*_1_. As a consequence, Eq. 1 may be inappropriate to describe phenomenon such as survival in the presence of natural selection, where competition for the same resource means that not all can survive. In some cases it may be possible to include a different model for the step or steps where Eq. 1 fails, analogously to the clonal expansion model [6] described in Section 6. But in principle, an alternative model may be required. We will return to this point in Section 9.

The rest of this section limits the discussion to situations where the paths to failure are independent, but where the failure-rate depends on the order of events. Important humanitarian examples are “cascading hazards” [19], where the risk of a disaster such as a mud slide is vastly increased if e.g. a wildfire occurs before it. An equivalent scenario would require *m* parts to fail for the system to fail, but the order in which the parts fail, modifies the probability of subsequent component failures. As an example, if three components A, B, and C, must fail, then we need to evaluate the probability of each of the 6 possible routes in turn, and obtain the overall failure probability from Eq. 1. Assuming the paths to failure are independent, then there are m! routes, giving 6 in this example. Writing the 6 routes as, 1=ABC, 2=ACB, 3=BAC, 4=BCA, 5=CAB, 6=CBA, and reading e.g. ABC as “A, then B, then C”, the survival probability is,

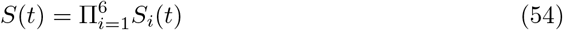

For failure by a particular route ABC we need the probability for the sequence of events, 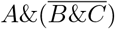, then 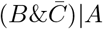, then *C*|(*AB*). We can calculate this using Eq. 16, for example giving,

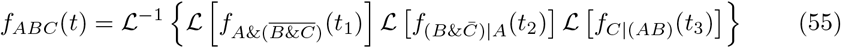

from which we can construct 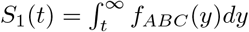.

Although in principle every term in e.g. Eqs. 54 and 55 need evaluating, there will be situations where results simplify. For example, if one route is much more probable than another - e.g. if it is approximately true that landslides only occur after deforestation, that may be due to fire, then we only need to evaluate the probability distribution for that route. As another example, if all the *f_i_* are exponentially distributed with different rates, then *f_ABC_* will be described by Moolgavkar’s solution. A more striking example is when there are very many potential routes to failure, as for the Armitage-Doll model where there are numerous stem cells that can cause cancer. In those cases, if the overall failure rate remains low, then the *f_i_*(*t*) in Eq. 55 must all be small with *S* ≃ 1 and *f* ≃ *h*, and can often be approximated by power laws. For that situation we have a general result that *f_i_*, *F_i_*, and *H_i_* will be a powers of time, and Eq. 2 gives,

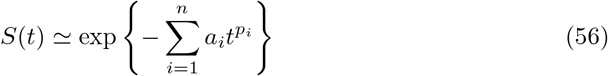

for some *a_i_* > 0 and *p_i_* > 0. Then *F*(*t*) = 1 − *S*(*t*), *f*(*t*) = −*dS*/*dt*, and *h*(*t*) ≃ *f*(*t*), can be approximated by a sum of power series in time. If one route is much more likely than the others then both *f*(*t*) and *h*(*t*) can be approximated as a single power of time, with the approximation best at early times, and a cross-over to different power-law behaviour at later times.

## 9 Cancer evolution, the tissue micro-environment, and model limitations

Cancer is increasingly viewed as an evolutionary process that is influenced by a combination of random and carcinogen-driven genetic and epigenetic changes [2, 3, 34, 33, 35, 36, 29, 21, 37], and an evolving tissue micro-environment [38, 39, 40, 41]. Although there is evidence that the number of stem cell divisions is more important for cancer risk than number of mutations [42, 43], the recognition that cells in a typical cancer are functionally and genetically diverse has helped explain cancers’ resistance to treatment, and is suggesting alternative strategies to tackle the disease through either adaptive therapies [44, 45, 46, 47] or by modifying the tissue’s micro-environment [48, 49, 39, 41]. This highlights two limitations of the multi-stage model described here.

### Evolution

As noted in Section 8, Eq. 1 cannot necessarily model a competitive process such as natural selection, where the growth of one cancer variant can inhibit the growth of another. If the process can be described through a series of rate-limiting steps, then we could still approximate it with a form of Eq. 16. Otherwise, the time-dependence of a step with competitive evolutionary processes may need to be modelled differently [30, 31], such as with a Wright-Fisher model [32, 31], or with an approximation such as the logistic model used to describe myeloid leukemia [6]. As emphasised by some authors [50, 39], a large proportion of genetic mutations occur before adulthood. Therefore it seems possible that some routes to cancer could be determined prior to adulthood, with genetic mutations and epigenetic changes in childhood either favouring or inhibiting the possible paths by which adult cancers could arise. If this led to a given cancer type occurring with a small number of sufficiently different incident rates, then it might be observable in a population’s incidence data as a mixture of distributions.

### Changing micro-environment

Another potential limitation of the model described in Section 5 is that the time to failure at each step is independent of the other failure times, and of the time at which that step becomes at risk. If the tissue microenvironment is changing with time, then this assumption fails, and the failure rate at each step is dependent on the present time. This prevents the factorisation of the Laplace transform used in Eqs. 13–15, that led to Eq. 16 for failure via *m* sequential steps. We can explore the influence of a changing micro-environment with a perturbative approximation. The simplest example is to allow the {*μ_j_*} in the Armitage-Doll model to depend linearly on the time 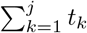 at which step *j* is at risk. Then the Armitage-Doll approximation of *f_j_*(*t_j_*) ≃ *μ_j_* for *μ_j_t_j_* ≪ 1, is replaced by

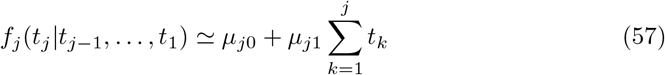

The calculation in Section 5 is modified, with,

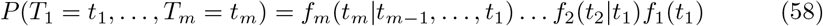

giving,

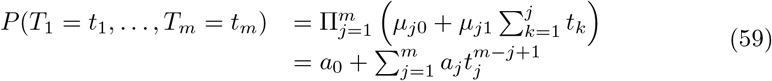

with 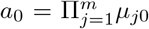, and {*a_j_*} being sums of products of *j* − 1 factors from {*μ*_*j*0_} and *m* − *j* + 1 factors from {*μ*_*k*_1__}. Replacing 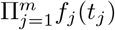 in Eqs. 13 and 14, with the right-side of Eq. 59, and evaluating the *m* integrals then gives,

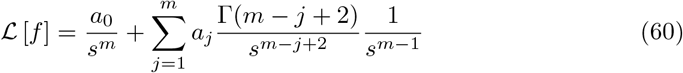

with solution,

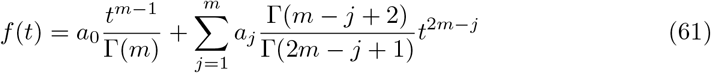

If the tissue micro-environment is changing rapidly enough that a term 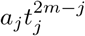 becomes comparable to or larger than *a*_0_*t*^*m*−1^, then the solution to Eq. 61 can behave like a larger power of time than the usual *m* − 1 for *m* rate-limiting steps. It is even possible for the incidence rate to slow or even *decrease*, if coefficients in Eq. 61 are negative. The example illustrates that if the micro-environment modifies cancer risk and is changing over a person’s lifetime, then it has the potential to strongly influence the observed rate of cancer incidence. The argument can be repeated with less generality or greater sophistication, e.g. expanding the coefficients *μ_i_* in the terms exp(−*μ_j_t_j_*) that appear in Moolgavkar’s model. Such models will have a complex relationship between their coefficients that might make them identifiable from cancer incidence data. This goes beyond the intended scope of this paper.

## 10 Conclusions

The purpose of this article is to provide a simple mathematical framework to describe existing multi-stage cancer models, that is easily adaptable to model events such as failure of complex systems, cascading disasters, and the onset of disease. The key formulae are Eqs. 1, 4, and 16 or equivalently 18, and a selection of analytical results are given to illustrate their use. Limitations of the multi-stage model are discussed in Sections 8 and 9. The examples in Section 6 can be combined in numerous ways to construct a wide range of models. Together the formulae are intended to provide a comprehensive toolkit for developing conceptual and quantitative models to describe failure, disaster, and disease.

## Acknowledgments

Thanks to Benjamin Cairns and Andrii Rozhok for helpful comments. This research was funded by a fellowship from the Nuffield Department of Population Health, University of Oxford, and with support from Cancer Research UK (grant no. C570/A16491).

## Supporting Information

### S1 Appendix Derivation of Eq. 23, and its relationship to Schwinger/Feynman parameterisations

The solution of Eq. 16 can be written in terms of multiple definite integrals that are sometimes easier to evaluate or approximate than directly evaluating Eq. 16. It is equivalent to expressing the solution as multiple convolutions using Eq. 21, and changing variables appropriately. The equation is obtained by Taylor expanding all functions before taking their Laplace transform, inverting the Laplace transform of the product of all terms (which is easy to do for the powers of time that appear in a Taylor expansion), then using a product of Beta functions to factorise and re-sum the resulting expression. In mathematical notation,

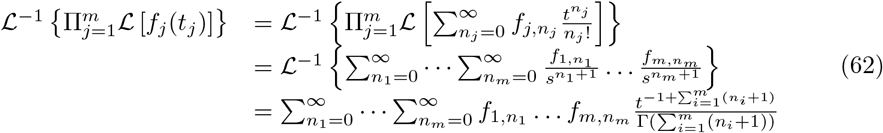

where *f_i,n_j__* = *∂^n_j_^f_i_*(*t_i_*)/*∂t*^*n_j_*^|_*t_i_*=0_. Now noting that the Beta function has,

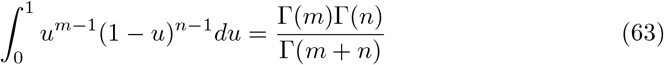

we can write,

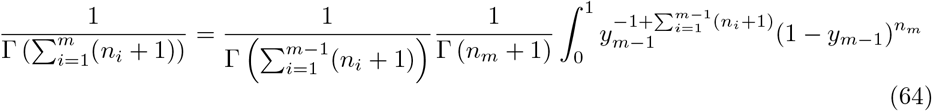

Repeatedly using this gives,

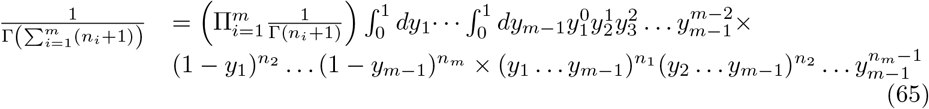

Using Eq. 65 to replace 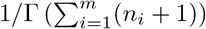 in Eq. 62, and grouping terms,

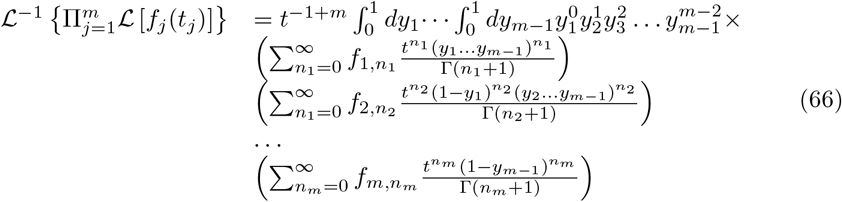

The *m* Taylor series can now be re-summed to give,

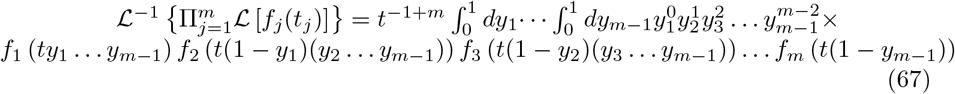

For example, taking *m* = 2 gives,

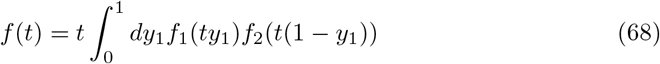

as we could have got from the convolution formula after a simple change of variables.

Eq. 67 might equivalently be regarded as a generalisation of a Schwinger/Feynman parameterisation, with,

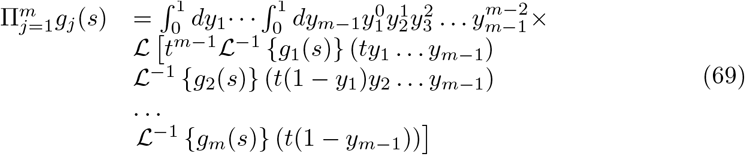

For example, taking *g_j_*(*s*) = 1/(*s*+*a_j_*)^*p_j_*^ and noting that 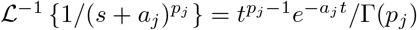, then we get,

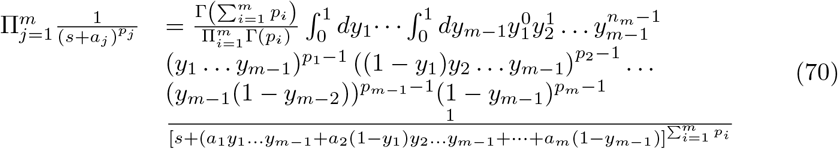

Taking *s* = 0, *m* = 2, and *p_j_* = 0 for all *j*, gives the most well-known form, with [27],

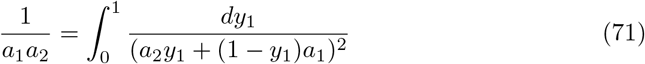

The identity Eq. 70 can be confirmed by writing the denominator as (*A_m_*)^*m*^, with,

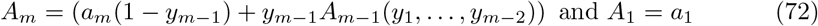

and integrating with respect to each of *y*_*m*−1_ to *y*_1_ in turn. For example, using the substitution *u* = *y*_*m*−1_/(1 + *α*_*m*−1_*y*_*m*−1_) with *α*_*m*−1_ = (*A*_*m*−1_ − *a_m_*)/*a_m_* and integrating between *u* = 0 and *u* = 1/(1 + *α*_*m*−1_), the integrand becomes (1/*a_m_*)(1/*A*_*m*−1_^*m*^). Repeating this for *y*_*m*−1_ to *y*_1_ confirms the identity.

## Notes

#### Summary of Updates

Final, pre-publication revisions.

